# Molecular bases and specificity behind the activation of the immune system OAS/RNAse L pathway by viral RNA

**DOI:** 10.1101/2024.07.08.602453

**Authors:** Emma Jung-Rodriguez, Florent Barbault, Emmanuelle Bignon, Antonio Monari

**Author notes:** Correspondence: A.M.

## Abstract

The first line of defense against invading pathogens usually relies on the innate immune systems. In this context the recognition of exogenous RNA structure is primordial to fight, notably, against RNA viruses. One of the most efficient immune response pathways is based on the sensing of RNA double helical motifs by the oligoadenylate synthase (OAS) proteins, which in turns triggers the activity of RNase L and, thus, cleaving cellular and viral RNA. In this contribution by using long range molecular dynamics simulation, complemented with enhanced sampling techniques, we elucidate the structural features leading to the activation of OAS by interaction with a model double strand RNA oligomer mimicking a viral RNA. We characterize the allosteric regulation induced by the nucleic acid leading to the population of the active form of the protein. Furthermore, we also identify the free energy profile connected to the active vs. inactive conformational transitions in presence and absence of RNA. Finally, the role of two RNA mutations, identified as able to down-regulate OAS activation, in shaping the protein/nucleic acid interface and the conformational landscape of OAS are also analyzed.

## 1. Introduction

RNA viruses are a class of pathogens which represent a significant threat due to their capability to induce epidemic or pandemic outbreak [1]. Indeed, the World Health Organization (WHO) has classed different RNA viruses along the most serious emerging infectious diseases [2]. The stunning evidence of the potential disruptive threat of RNA virus has come to light with the emergence of the COVID-19 pandemic outbreak [3–7], which started at the end of 2019 in Wuhan China and spread to virtually every continent by the spring of 2020, pushing authorities to implement severe social distance measures, including full lockdown. COVID-19 infection is caused by a β-stranded positive RNA virus belonging to the *coronaviridae* family [3], which has been named SARS-CoV-2, and shares homology with other potentially lethal coronaviruses such as SARS-CoV [8], appeared in 2003, and MERS-CoV, emerged in 2013 [9]. The RNA viruses encompass other extremely virulent pathogens, such as hemorrhagic fever viruses (Ebola), flaviruses and arboviruses including Zika [10,11], West Nile [12], and Chikungunya viruses [13]. Interestingly, since the latter pathogens exploit insect, particularly mosquitos, as vectors, their worldwide diffusion has increased significantly as a consequence of global warming [14], which allows the colonization of temperate regions by vectors originally developing in tropical areas. Furthermore, the common seasonal influenza, which is, however, each year correlated with a non-negligible mortality rate and has been at the origin of serios pandemics events in the XX^th^ century, is also caused by a rapidly mutating genome-segmented RNA virus [15,16].

The frontline of the immune system defense against the invasion by RNA in vertebrates usually relies on the innate immune system and in particular on its capacity to sense the presence of exogenous genetic material, such as specific RNA sequences and structural motifs [17,18]. The antiviral innate immune response is usually characterized by the activation of pro-inflammatory signaling, as well as the production of interferon and cytokine [19], thus, developing an unfit microenvironment and inducing the apoptosis or senescence of infected cells. The combination of these responses is aimed at slowing the viral spreading limiting its diffusion. Different innate immune systems pathways have been characterized in the last years. These involve the cGAS/STING pathway [20], which is activated by the presence of exogenous DNA, and in a less extent RNA [21], and is mediated by the signaling exerted by small cyclic nucleotides. Interestingly, the overactivation of the STING protein has been correlated to the insurgence of serious COVID-19 outcomes, and in particular to the cytokine storm which is one of the main causes of morbidity of SARS-CoV-2 [22]. Yet, the main reaction to the presence of pathogen RNA is the one mediated by the 2’, 5’-oligoadenylate synthase (OAS) and the RNAse L proteins [23–25]. In this pathway the catalytic activity of OAS is triggered by its interaction with double strand RNA fragments, and leads to the production of short 2′-5′-oligoadenylate oligomers. The latter in turn are used as signal to mediate the further activation of the RNAse L endonu-cleases which cleaves, non-specifically, single strand RNA. The cleavage of both viral and cellular RNA, including messenger RNA, leads to the apoptosis of the cell, and thus, stops the viral reproduction and diffusion [24]. The role and the efficiency of the OAS pathway in the defense against viral aggression has been particularly underlined, recently, in the case of SARS-CoV-2, in which human haplotypes of Neanderthal descendance which preferentially colocalize with the reproduction compartment of the virus, have been correlated to milder or even symptomatic infections [26].

Usually, three main variants of OAS are encountered in vertebrates [27]. The smaller OAS1 presents only one catalytic unit and is particularly efficient in recognize short double-strand RNA oligomers [28]. Two catalytic cores are bridged together in the case of the OAS2 variant, leading to the recognition of medium-length RNA oligomers, while three units are bridged in the case of OAS3, thus allowing the interaction with longer strands. Interestingly, only two of the catalytic centers in OAS3 are active, while the third is not reactive but participate to the recognition of viral RNA.

From a biochemical and biophysical point of view, as shown in Figure 1 for OAS1 [29–31], the binding of viral RNA involves a large recognition area which is spatially distant from the catalytic centers, which presents a [Mg_2_]^4+^ cluster, similarly to the majority of polymerases and nuclease enzymes. Therefore, the activation of OAS should involve an RNA-driven allosteric regulation of the catalytic activity, which allows the activation of the pathway only in presence of exogenous genetic material. Recently, it has been shown that the human OAS1 presents a strong selectivity toward the 5’-untranslated region (5’-UTR) of SARS-CoV-2 genome, and in particular its first stem loop motif (SL1) [32,33]. The affinity of OAS for this motif could also explain its specific activation since 5’-UTR, and SL1 in particular, are highly conserved regions and exert an important regulatory role in the viral cycle, in particular concerning RNA translation and replication [34,35]. Similar structural regions involving stem loops and presenting regulatory activity are also found in other RNA viruses, such as Dengue [36], West Nile [37], or Zika [38], and are specifically recognized by OAS1 [39]. By using long-range classical molecular dynamic (MD) simulations we have shown that the while the specific recognition of SARS-CoV-2 SL1 is mainly driven by sequence specific interactions with the extruded nucleobases [32], the tertiary arrangement of the West Nile SL1 offers structural-based specific interaction patterns [39]. Crystal structure of OAS1 in active form [29], i.e. complexed with model double stranded RNA (in the following referred as holo structure) has been obtained, as well as its inactive counterpart in absence of RNA (apo form), revealing important structural differences. In addition, as shown by Donovan et al. [29] site-specific mutations of the RNA sequence down-regulate the activation of OAS1. Yet, the molecular and atomistic factors dictating the allosteric regulation of OAS1 remains partially elusive. Therefore, in this contribution we scrutinize the human OAS1 in active and inactive form and the interplay with RNA in modulating the allosteric conformational transition of OAS1. In addition, we also analyze the structural effects of RNA point mutations [29] and the effects of RNA in modulating the free energy profile of the active vs. inactive transition of OAS1.

**Figure 1.**
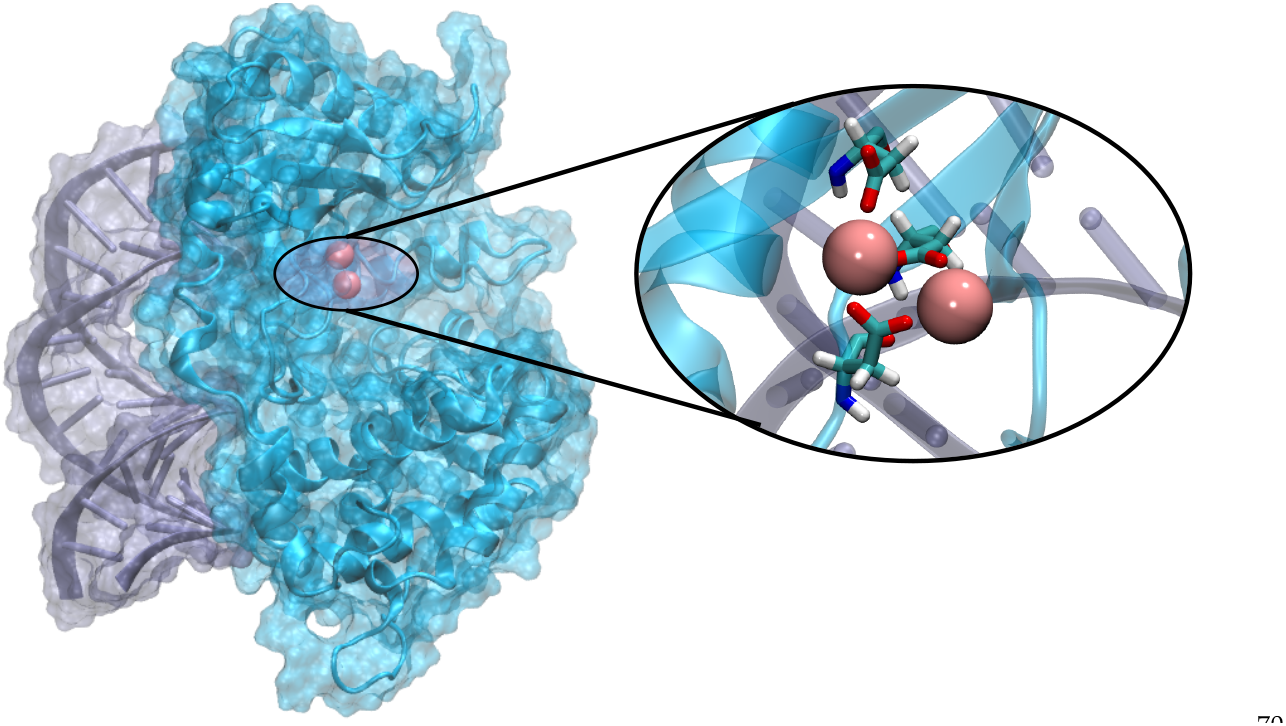
Representative snapshot showing the structure of the human OAS1 protein (in cyan) interacting with a model RNA double strand (in purple). In the inlay a zoom of the active site showing the Mg_2_^4+^ cluster complexed by aspartate ligands.

## 2. Materials and Methods

The initial structure of the human OAS1 enzyme in active form has been retrieved from the PDB data base (id 4IG8) [29]. The structure also included the presence of a 18-mer RNA double strand of sequence 5’-GGCUUUUGACCUUUAUGC-3’ which has been maintained (HOLO system). Missing residues have been added using the Swiss-Model online server [40]. No structure of the human OAS in inactive apo form could be found in the pdb database. Therefore, we have constructed an Apo form of OAS1 simply removing the RNA double-strand from the structure of PDB 4IG8, this system will be further referred to as APO1. In addition, we have constructed a second model (APO2) using the crystal structure of the porcine OAS1 in its apo form (PDB id: 4RWQ) [25], which shares a 74% homology with the human protein, as a template for homology model, using again the Swiss-model server. To simulate the effects of the mutation of the RNA in the affinity towards OAS1 we also constructed two further systems by manually mutating the base pairs 17 (GC17AU) and 18 (GC18AU), referred to as HOLO17 and HOLO18 in the following, respectively.

All the systems have been soaked in a cubic water box enforcing a 9 Å buffer, using the amber tleap utilities [41] and neutralized with the minimum amount of K^+^ ions. The OAS protein has been modeled using the amberff19SB force field [42], while RNA is described with OL3 RNA force field [43] and solvent, water, with the TIP3P model [44,45]. All the equilibrium MD simulations have been performed in the isotherm and isobaric (NPT) ensemble at a temperature of 300 K and pressure of 1 atm, which have been enforced using the Langevin thermostat [46]and barostat [47]. Particle Mesh Ewald (PME) summation with a cut-off of 9 Å have been consistently used. The Hydrogen Mass Repartition (HMR) strategy [48] has been used in combination with Rattle and Shake algorithms [49]to allow using a time step of 4 fs for integrating the Newton equations of motion. Prior to the production run the system has been optimized using conjugated gradient algorithm and equilibrated and then thermalized by progressively removing constraints on the RNA and protein backbone atoms during three consecutive steps of 36 ns each. All the MD simulations have been performed using the NAMD code [50,51], and analyzed with VMD [52] and cpptraj [53] utilities.

In addition to equilibrium MD simulations, and to enforce the transition between the active and inactive conformations in the HOLO and APO systems, we have performed enhanced sampling MD simulations. To this aim the difference in the backbone root mean square distribution (ΔRMSD) between representative structures of the two conformations, obtained from the equilibrium MD, has been used as the collective variable. The free energy profile connecting the two conformations has been obtained by Umbrella Sampling (US) procedure using forces comprised between 1.0 and 10.0 N. The suitability of the collective variable has been previously examined performing Steered Molecular Dynamics (SMD) assuring that the path is indeed connecting the two equilibrium conformations. Enhanced sampling has been performed using NAMD coupled to the Colvar [54] utilities. The cavity has been identified and quantified using the CASTpFold webserver [55].

## 3. Results

The MD simulations of both the HOLO and the reconstructed APO1 and APO2 models are stable and converged at the μs time-scale as can be observed notably from the analysis of the time-evolution of the RMSD, which is reported in SI. In Figure 2 we report the superposition of the structures obtained for both APO models with the HOLO system. Of note, in presence of double strand RNA, and at least at the time scale of our dynamics, the HOLO system experiences a remarkable conformational stability, presenting only very limited deviation from the crystal structure. As shown in Figure 2A the APO1 structure which has been constructed by simply removing the RNA double strand is not able to recover the native structure and remains locked in a conformation which presents an almost perfect superposition with the HOLO system. On the contrary, significant structural changes can be observed in the case of the APO2 system. This concerns in particular the position of the two α-helices involving residues 86-110 and 201-220, respectively and the sliding and rotation of the β -sheet region. Interestingly, the most important global structural changes do not involve areas which are in direct contact with the RNA recognition interface, thus confirming a halosteric regulation of the activation of OAS1. Indeed, the changes observed may instead affect the accessibility of the active site, involving the Magnesium cluster catalytic center. This effect can be directly assessed by measuring the volume of the cavity encompassing the active center, which amounts at 2.6 and 2.0 nm^3^ for HOLO and APO2. Interestingly, and confirming the locking of this conformation in a metastable state, APO1 presents a value of the cavity volume absolutely comparable with the one of the original HOLO system. Our MD simulations are also in line with the results inferred by Donovan et al. [29] by analyzing the different crystal structures. Our results are then coherent in identifying an active conformation, which is populated in the HOLO structure upon interaction with RNA, and an inactive state which is characteristic of OAS1 at rest. In this respect, we may consider that APO1 should not be considered as a physically or biologically relevant state, but instead as a computational artefact.

**Figure 2.**
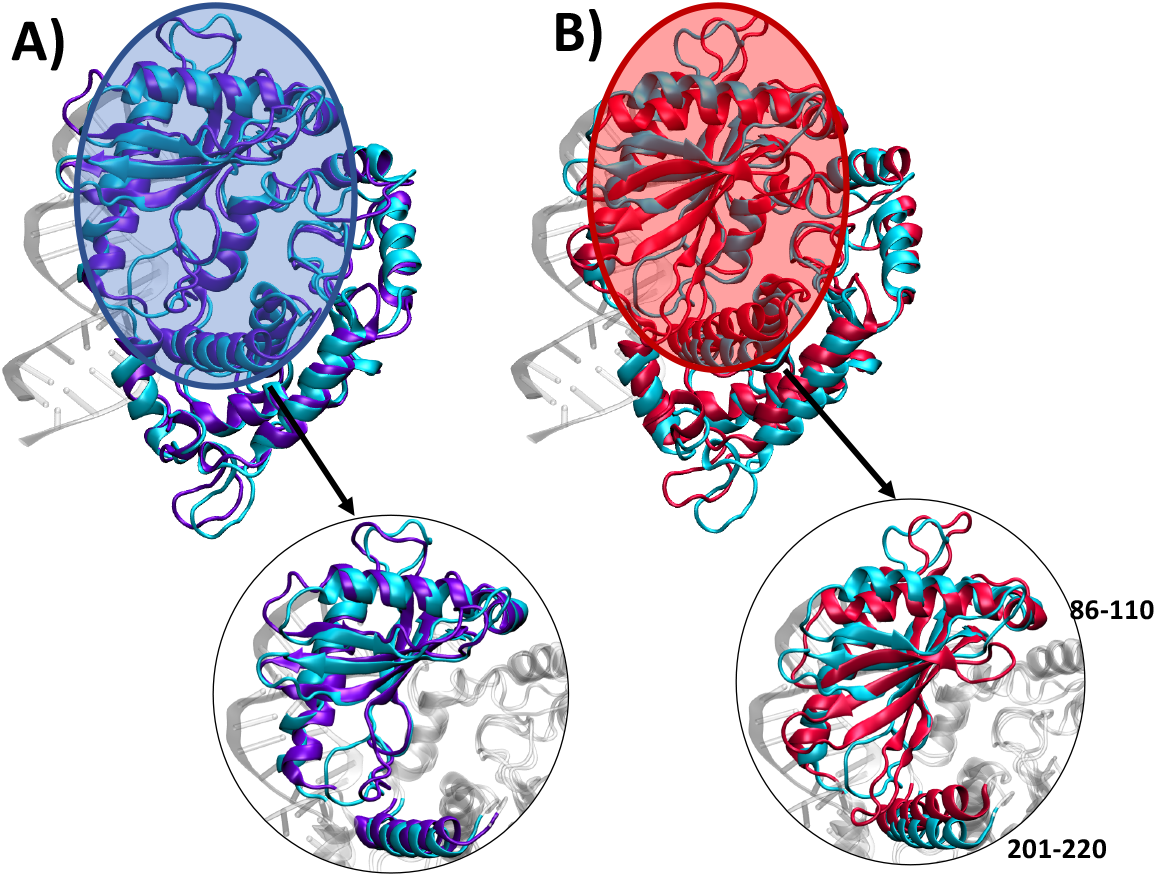
Superposition of representative snapshots issued from the MD simulation between the HOLO (cyan) and the APO1 (panel A in purple) and APO2 (panel B in red) A zoom of the region mostly affected by the conformational changes is also provided.

When switching to the analysis of the RNA-induced structural reorganization at a residue level, it was pointed out [29] that the allosteric propagation towards the active site was mainly driven by the basic residues K66 and R195 which should switch their position to optimize their interactions with E233.

As can be appreciated from Figure 3A the transition between the HOLO and APO2 form involves a rather important rearrangement of this triad, in particular concerning the placement of K66. On the contrary as reported in SI no significant reorganization can be observed for APO1, once again pointing to the fact that this conformation is locked in a metastable state. Interestingly, from the time evolution of the residue-residue distance reported in Figure 3B it can be appreciated that while in APO conformation K66 is isolated in the HOLO conformation it develops a stable interaction with both the side chain of E223 and the backbone of R195. However, no direct interaction between R195 and E223 can be observed in neither HOLO or APO simulations. Therefore, our results may suggest that, differently from what was supposed from the analysis of the different crystal structures, the conformational transition between the active and inactive form does not involve a switching between the positions of the two basic residues, but rather a double role of K66 which bridges together E223 and R195, exploiting both electrostatic interactions with the side chains, and hydrogen bond stabilization with the backbone. This observation is also confirmed considering the average distances between the centers of mass of the K66 and E223 residues in HOLO and APO conformations which amounts to 5.8 ± 1.0 and 17.1 ± 1.5 Å, respectively. While the average distance of the center of mass between K66 and R195 is of 9.0 ± 1.0 and 23.7 ± 2.0 Å, for the HOLO and APO2 conformations, respectively. The locking of K66 in the HOLO form may also be inferred from the variation of the root mean square fluctuation reported in SI (Figure S8 and S9) which shows the strong reduction of the flexibility of this residue upon recognition of the RNA.

**Figure 3.**
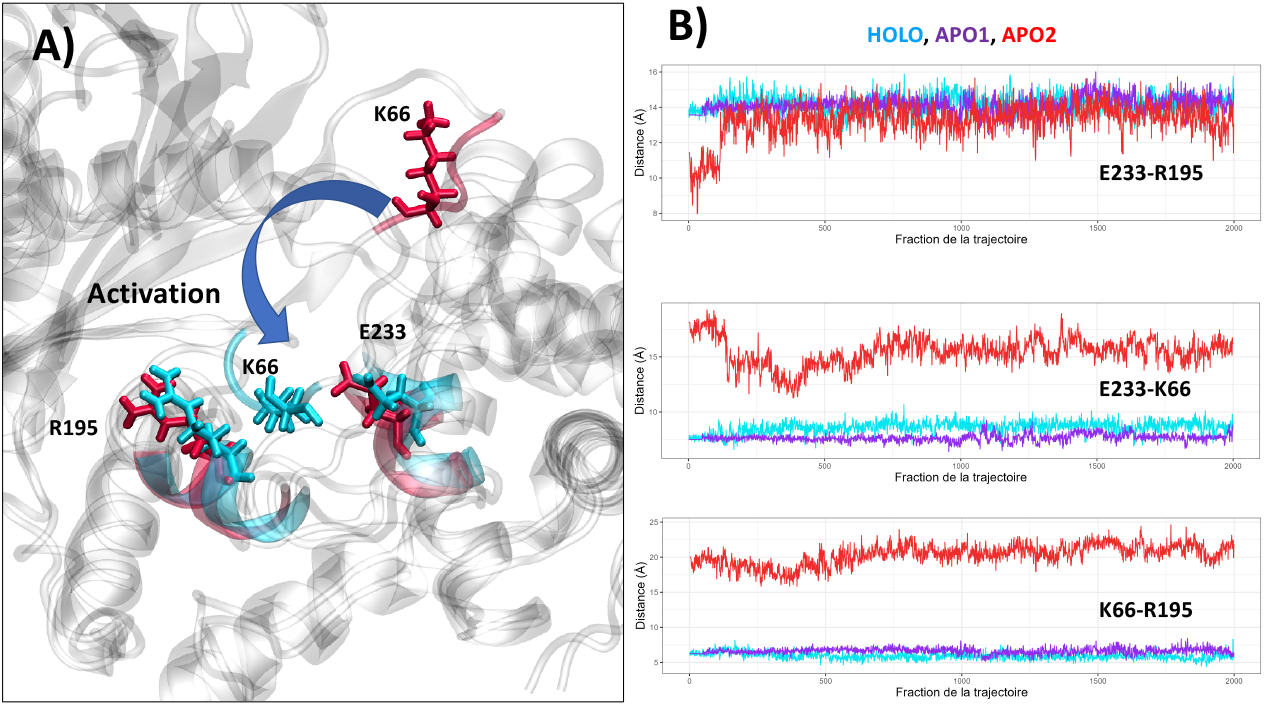
A) Superposition of the HOLO (cyan) and APO2 (red) structures highlighting the position of the residues E223, K66, and R195 in the two structures. B) Time evolution of the distances between the residues E233, R195, and K66 in HOLO (cyan), APO1 (purple), and APO2 (red).

To better assess the thermodynamic and kinetics of the conformational transitions provoked by the interaction with RNA we have used umbrella sampling enhanced MD simulations to enforce the active/inactive conformational transition in HOLO and APO systems, i.e. in presence and in absence of RNA. The free energy profiles related to this transition, along the variation of the RMSD between the two equilibrium states were generated, clarifying the thermodynamics behind OAS1 activation.

The free energy profile and potential of mean force ((PMF) are reported in Figure 4., from which it can be readily observed that the global minimum of the APO2 system corresponds to the inactive conformation (ΔRMSD = -2.6 Å). However, in the case of the HOLO system the active conformation (ΔRMSD = +2.6 Å) is more favorable. Interestingly, the free energy increase when pushing the active conformation of the HOLO form (with RNA) towards the inactive one corresponds to about +12 kcal/mol. Conversely, the penalty of maintaining an active conformation in absence of RNA amounts to about 10 kcal/mol. In both HOLO and APO systems, the transition towards the most stable conformation proceeds rather smoothly, without the need to bypass significant free energy barriers (the highest one amounting to 2 kcal/mol). This result is coherent with the biological role of OAS1, which should be rapidly activated upon sensing the presence of viral RNA, and should be readily inactivated in absence of exogenous RNA. On the other hand, the fact that the inactive conformation was not recovered for the APO1 system should probably be due to the large inertia correlated to the extended conformational rearrangement. As a matter of fact, an equilibrium MD simulation in which the RNA double strand was manually docked onto APO2 system in inactive conformation has shown a partial recovering of the active form (see ESI).

**Figure 4.**
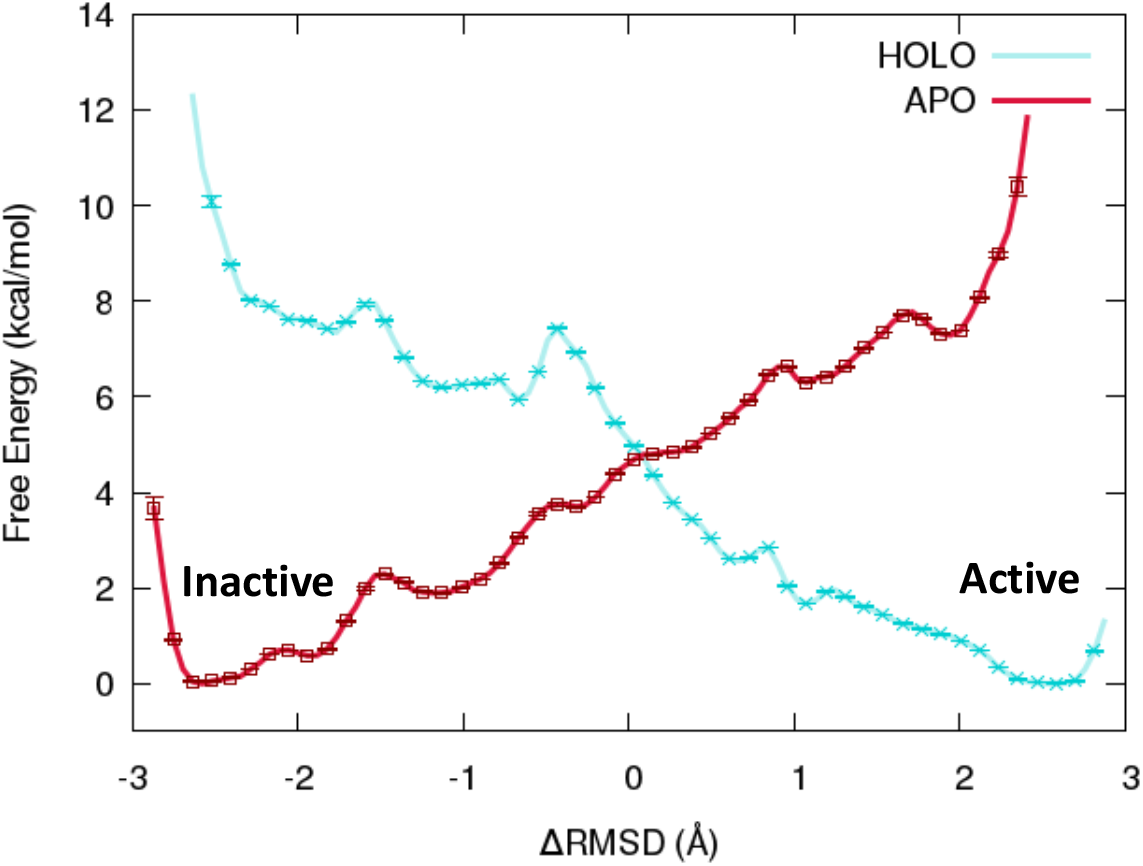
Potential of mean force for the active/inactive transition in HOLO (cyan) and APO2 (red) systems obtained from umbrella sampling enhanced MD simulations.

After considering the thermodynamics and kinetic factors determining the conformational equilibrium between active and inactive form upon recruitment or release of double strand RNA oligomers, we scrutinized more into details the inherent factors stabilizing the protein/nucleic acid interface, and hence favoring recognition. In Figure 5A we report a representative snapshot highlighting the interacting interface between OAS1 and our model DNA. Unsurprisingly, the interface presents a high density of basic, positively charged residues, mainly lysine and arginine. This accumulation of basic residues participates in creating an electrostatically positive channel which may efficiently accommodate the negatively charged RNA backbone. The presence of such a recognition motif is also widely common in proteins interacting with nucleic acid, including histones and polymerases. While the electrostatic interface clearly represents the main factor maintaining the RNA/OAS1 complex, it is important to underline that other non-covalent interactions take place and contribute to the stabilization of the interface. Interestingly, as shown in Figure 5B we may also identify a cluster of hydrophobic residues at the center of the RNA/protein interface. These residues may reinforce the specificity of the recognition by developing favorable interactions with the nucleic acid nucleobases. Indeed, it has also been recently shown that in some cases the recognition of viral material by OAS1 may also be mediated by the interaction with extruded nucleobases [32].

**Figure 5.**
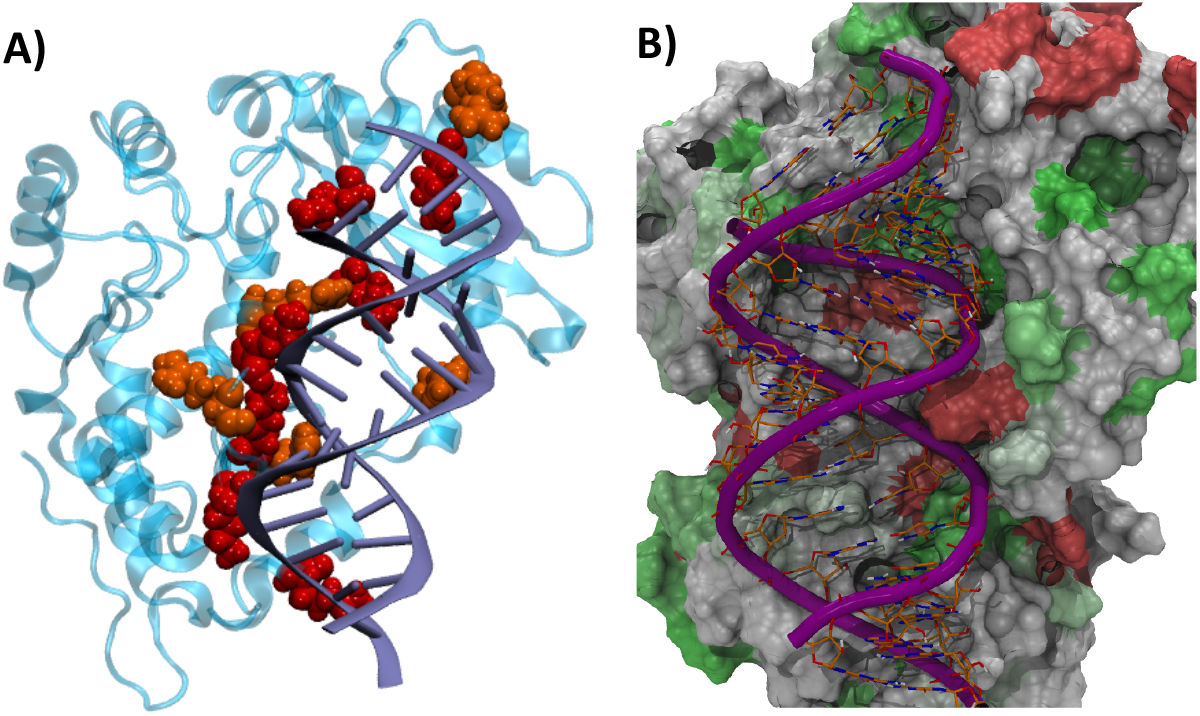
A) The interaction interface between OAS1 and the model double-strand RNA model. The protein interfacial basic residues (lysine red and arginine orange) are highlighted in van der Waals representation. B) Representation of the hydrophobicity of the OAS1 protein in interaction with RNA. The protein is rendered in surface representation. Residues colored in green are hydrophobic, in red hydrophilic, and in white amphiphilic.

In particular the analysis of the MD simulation for the HOLO system has shown the establishment of some rather persistent hydrogen bonds involving also the peripheral nucleobases and the nearby protein residues. The summary of the different hydrogen bonds, together with their persistence along the MD simulation are reported in Table 1. It is evident that in the original system the RNA/OAS1 interface is also stabilized by persistent hydrogen bonds, which are also characterized by near ideal angle and distances between the donor and the acceptor. These results are also coherent with previous MD simulations devoted to the analysis of the specific recognition of SARS-CoV-2 5’-UTR domain, which has shown the sequence-dependent additional stabilization by hydrogen bonds with the extruded nucleobases [32].

**Table 1.**
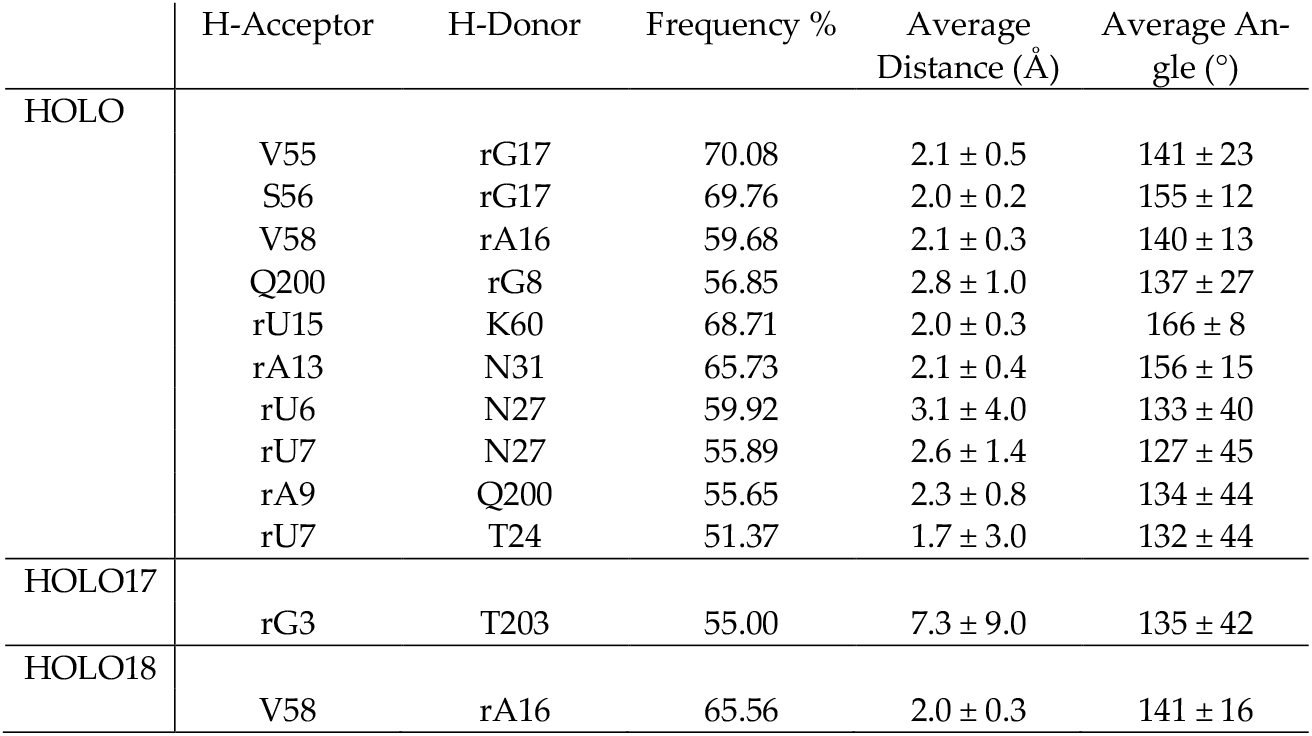
Main hydrogen bond interactions taking place along the MD simulation for the native HOLO and the mutated HOLO17 and HOLO18. The frequency of occurrence is given together with the average distance and angle calculated along the MD simulation. Residues belonging to RNA are preceded by an uncapitalized ‘r’ for clarity.

|  | H-Acceptor | H-Donor | Frequency % | Average Distance (Å) | Average Angle (°) |
| --- | --- | --- | --- | --- | --- |
| HOLO | V55 | rG17 | 70.08 | $2.1 \pm 0.5$ | $141 \pm 23$ |
| | S56 | rG17 | 69.76 | $2.0 \pm 0.2$ | $155 \pm 12$ |
| | V58 | rA16 | 59.68 | $2.1 \pm 0.3$ | $140 \pm 13$ |
| | Q200 | rG8 | 56.85 | $2.8 \pm 1.0$ | $137 \pm 27$ |
| | rU15 | K60 | 68.71 | $2.0 \pm 0.3$ | $166 \pm 8$ |
| | rA13 | N31 | 65.73 | $2.1 \pm 0.4$ | $156 \pm 15$ |
| | rU6 | N27 | 59.92 | $3.1 \pm 4.0$ | $133 \pm 40$ |
| | rU7 | N27 | 55.89 | $2.6 \pm 1.4$ | $127 \pm 45$ |
| | rA9 | Q200 | 55.65 | $2.3 \pm 0.8$ | $134 \pm 44$ |
| | rU7 | T24 | 51.37 | $1.7 \pm 3.0$ | $132 \pm 44$ |
| HOLO17 | rG3 | T203 | 55.00 | $7.3 \pm 9.0$ | $135 \pm 42$ |
| HOLO18 | V58 | rA16 | 65.56 | $2.0 \pm 0.3$ | $141 \pm 16$ |

It has been recently shown that two specific mutations in the double-strand RNA sequence, namely the GC17AU (HOLO17) and the GC18AU (HOLO18) correlates with a downregulation of the OAS1 activation [29]. From the results reported in Table 1 it appears evident that the hydrogen-bond network in the case of the mutated oligomers has been almost completely disrupted, with only one, relatively weak, hydrogen bond persisting for both mutants. Interestingly the hydrogen bond interactions have been lost also in case of nucleobases not directly involved in the two mutations, suggesting a subtly regulated tuning of the interfacial recognition. However, the sole observation of the native HB network perturbation masks a completely different structural organization between the two structures. Indeed, as shown in Figure 6, the recognition interface is completely lost in HOLO17. Instead, in the case of HOLO18, and despite the loss of the hydrogen bond interactions, the protein/RNA interface is relatively well conserved, even if the activation of OAS1 catalytic activity is reduced. These evidences further confirm that OAS1 is indeed able to enforce a more specific recognition of RNA sequences, thanks notably to the establishment of specific hydrogen bond networks. This is again coherent with the specific activation of the OAS/RNAse L pathway only in presence of exogenous infections, and thus avoid an overactivation of the innate immune system. It is also in line with our results evidencing the role of hydrogen bonding in leveraging the specific recognition of SARS-CoV-2 genetic material [32].

**Figure 6.**
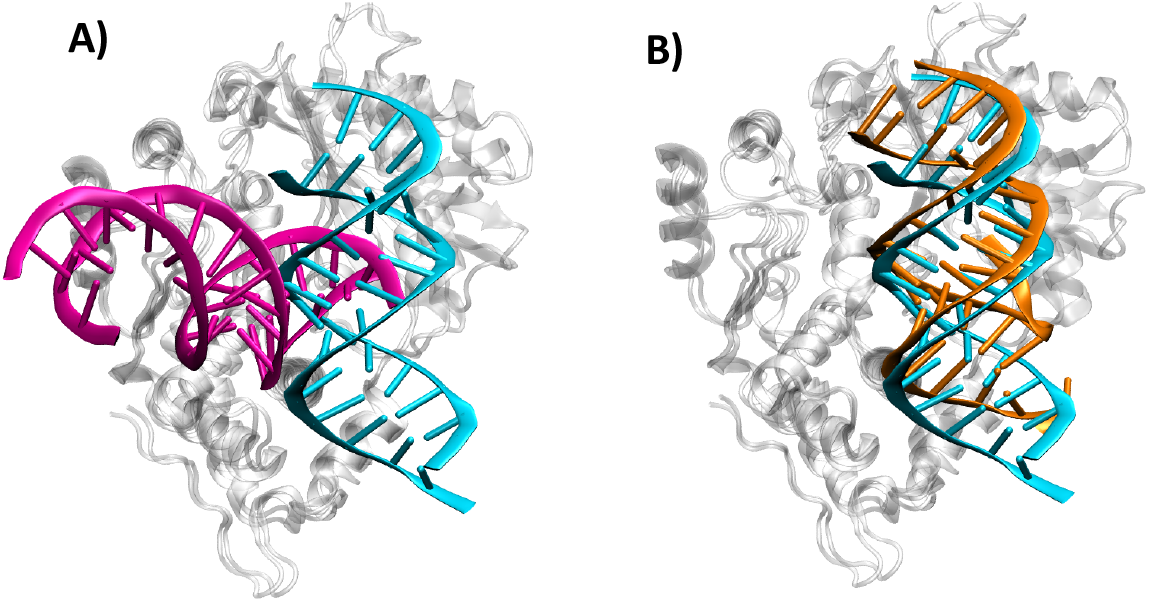
Representative structure of the HOLO17 (A, magenta) and HOLO18 (B, orange) mutants in complex with OAS1 and superposed to the original HOLO structure (in cyan).

## 4. Discussion and Conclusion

By using long-range MD simulations complemented with enhanced sampling and free energy methods, we have unraveled the molecular bases related to the subtle conformational equilibrium of OAS1 upon binding to double stranded RNA. Indeed, we have confirmed the presence of an active and an inactive conformation, which differ by the relative orientation of α-helixes and β-sheets. The allosteric regulation, leveraging the population of the active conformation, leads to an enlargement of the catalytic site pocket. In addition, we have shown that the driving force for the population of the active conformation is of about 12 kcal/mol and that the transition does not involve significant energy barriers. As concerns the RNA/protein interaction we have shown that, in addition to the expected salt bridges and electrostatic interactions, an extended network of hydrogen bonds is present that stabilizes the interface and drives the transition towards the active form. Indeed, we have shown that in the case of activity loss mutations involving the peripheral nucleobases of the double stranded oligomer, the native hydrogen bonds pattern is almost entirely disrupted. Interestingly, this situation may also lead to the complete loss of the correct RNA positioning in case of the GC17AU mutant. Our results, while confirming experimental observations and previous MD simulations, shed a further light on the thermodynamics driving the activation of OAS1, and thus on the triggering of the innate immune system upon recognition of exogenous genetic material. The role of hydrogen bonds, which leads to sequence specific recognition of the RNA strands and adds up to the structural recognition, has also been clearly evidenced. Such a knowledge can help in understanding the factors tuning the immune system response, and eventually in designing specific immunotherapy drugs aimed against viral or autoimmune diseases.

In the future we plan to extend this study elucidating the thermodynamics and inactive/active transition of OAS1 taking into account its interaction with actual viral sequences such as the first stem loop in 54-UTR of SARS-CoV-2 or West Nile viruses.

## Supplementary Materials

The following supporting information can be downloaded at: www.mdpi.com/xxx/s1, Details on the definition of the collective variable. Time evolution of the RMSD for the different systems studied 5figure S1-S4, S6, S7) Superposition of the APO2+RNA and HOLO system (Figure S5). RMSF for the HOLO and APO2 systems (Figure S8) and their difference (S9). Distribution of the values of the collective variables over the umbrella sampling (Figure S10).

## Author Contributions

E.J.-R.; data acquisition, methodology. E.B., F.B. A.M.; conceptualization, funding acquisition, supervision. A.M.; original draft preparation. E.J.-R., E.B., F.B., A.M.; writing, review, and editing. All authors have read and agreed to the published version of the manuscript.

## Data Availability Statement

Data are available upon reasonable request.

## Acknowledgments

The authors thank GENCI and Explor computing centers for computational resources. ANR and CGI are acknowledged for their financial support of this work through Labex SEAM ANR 11 LABX 086, ANR 11 IDEX 05 02 The support of the IdEx “Université Paris 2019” ANR-18-IDEX-0001 and of the Platform P3MB is also gratefully acknowledged.

## References

1. Garcia-Blanco, M.A.; Ooi, E.E.; Sessions, O.M. RNA Viruses, Pandemics and Anticipatory Preparedness. Viruses 2022, 14, 2176.

2. Chakrabartty, I.; Khan, M.; Mahanta, S.; Chopra, H.; Dhawan, M.; Choudhary, O.P.; Bibi, S.; Mohanta, Y.K.; Emran, T. Bin Comparative overview of emerging RNA viruses: Epidemiology, pathogenesis, diagnosis and current treatment. Ann. Med. Surg. 2022, 79, 103985.

3. Astuti, I.; Ysrafil Severe Acute Respiratory Syndrome Coronavirus 2 (SARS-CoV-2): An overview of viral structure and host response. Diabetes Metab. Syndr. Clin. Res. Rev. 2020, 14, 407–412.

4. Tchicaya, A.; Lorentz, N.; Leduc, K.; de Lanchy, G. COVID-19 mortality with regard to healthcare services availability, health risks, and socio-spatial factors at department level in France: A spatial cross-sectional analysis. PLoS One 2021, 16, e0256857.

5. Brant, A.C.; Tian, W.; Majerciak, V.; Yang, W.; Zheng, Z.M. SARS-CoV-2: from its discovery to genome structure, transcription, and replication. Cell Biosci. 2021, 11.

6. Del Rio, C.; Omer, S.B.; Malani, P.N. Winter of Omicron-The Evolving COVID-19 Pandemic. JAMA 2021, 327, 319–320.

7. Watkins, J. Preventing a covid-19 pandemic. BMJ 2020, 368, m810.

8. Fehr, A.R.; Perlman, S. Coronaviruses: An overview of their replication and pathogenesis. In Coronaviruses: Methods and Protocols; Maier, H.J., Bickerton, E., Britton, P., Eds.; Springer New York: New York, NY, 2015; pp. 1–23 ISBN 9781493924387.

9. Rabaan, A.A.; Al-Ahmed, S.H.; Sah, R.; Alqumber, M.A.; Haque, S.; Patel, S.K.; Pathak, M.; Tiwari, R.; Yatoo, M.I.; Haq, A.U.; et al. MERS-CoV: epidemiology, molecular dynamics, therapeutics, and future challenges. Ann. Clin. Microbiol. Antimicrob. 2021, 20, 8.

10. Barrows, N.J.; Campos, R.K.; Liao, K.C.; Prasanth, K.R.; Soto-Acosta, R.; Yeh, S.C.; Schott-Lerner, G.; Pompon, J.; Sessions, O.M.; Bradrick, S.S.; et al. Biochemistry and Molecular Biology of Flaviviruses. Chem. Rev. 2018, 118, 4448–4482.

11. Kazmi, S.S.; Ali, W.; Bibi, N.; Nouroz, F. A review on Zika virus outbreak, epidemiology, transmission and infection dynamics. J. Biol. Res. 2020, 27.

12. Richter, J.; Tryfonos, C.; Tourvas, A.; Floridou, D.; Paphitou, N.I.; Christodoulou, C. Complete genome sequence of West Nile Virus (WNV) from the first human case of neuroinvasive WNV infection in Cyprus. Genome Announc. 2017, 5.

13. Oo, A.; Rausalu, K.; Merits, A.; Higgs, S.; Vanlandingham, D.; Bakar, S.A.; Zandi, K. Deciphering the potential of baicalin as an antiviral agent for Chikungunya virus infection. Antiviral Res. 2018, 150, 101–111.

14. Zell, R.; Krumbholz, A.; Wutzler, P. Impact of global warming on viral diseases: what is the evidence? Curr. Opin. Biotechnol. 2008, 19, 652–660.

15. Krammer, F.; Smith, G.J.D.; Fouchier, R.A.M.; Peiris, M.; Kedzierska, K.; Doherty, P.C.; Palese, P.; Shaw, M.L.; Treanor, J.; Webster, R.G.; et al. Influenza. Nat. Rev. Dis. Prim. 2018, 4, 3.

16. Javanian, M.; Barary, M.; Ghebrehewet, S.; Koppolu, V.; Vasigala, V.K.R.; Ebrahimpour, S. A brief review of influenza virus infection. J. Med. Virol. 2021, 93, 4638–4646.

17. Janeway, C.A.; Medzhitov, R. Innate immune recognition. Annu. Rev. Immunol. 2002, 20, 197–216.

18. Marshall, J.S.; Warrington, R.; Watson, W.; Kim, H.L. An introduction to immunology and immunopathology. Allergy, Asthma Clin. Immunol. 2018, 14.

19. Hornung, V.; Hartmann, R.; Ablasser, A.; Hopfner, K.P. OAS proteins and cGAS: Unifying concepts in sensing and responding to cytosolic nucleic acids. Nat. Rev. Immunol. 2014, 14, 521–528.

20. Bhat, N.; Fitzgerald, K.A. Recognition of cytosolic DNA by cGAS and other STING-dependent sensors. Eur. J. Immunol. 2014, 44, 634–640.

21. Webb, L.G.; Fernandez-Sesma, A. RNA viruses and the cGAS-STING pathway: reframing our understanding of innate immune sensing. Curr. Opin. Virol. 2022, 53.

22. Berthelot, J.M.; Lioté, F.; Maugars, Y.; Sibilia, J. Lymphocyte Changes in Severe COVID-19: Delayed Over-Activation of STING? Front. Immunol. 2020, 11, 607069.

23. Choi, U.Y. un.; Kang, J.S.; Hwang, Y.S. ahn.; Kim, Y.J. Oligoadenylate synthase-like (OASL) proteins: dual functions and associations with diseases. Exp. Mol. Med. 2015, 47, e144.

24. Schwartz, S.L.; Conn, G.L. RNA regulation of the antiviral protein 2′-5′-oligoadenylate synthetase. Wiley Interdiscip. Rev. RNA 2019, 10, e1534.

25. Lohöfener, J.; Steinke, N.; Kay-Fedorov, P.; Baruch, P.; Nikulin, A.; Tishchenko, S.; Manstein, D.J.; Fedorov, R. The activation mechanism of 2′-5′-oligoadenylate synthetase gives new insights into OAS/cGAS triggers of innate immunity. Structure 2015, 23, 851–862.

26. Di Maria, E.; Latini, A.; Borgiani, P.; Novelli, G. Genetic variants of the human host influencing the coronavirus-associated phenotypes (SARS, MERS and COVID-19): Rapid systematic review and field synopsis. Hum. Genomics 2020, 14, 1–19.

27. Leisching, G.; Cole, V.; Ali, A.T.; Baker, B. OAS1, OAS2 and OAS3 restrict intracellular M. tb replication and enhance cytokine secretion. Int. J. Infect. Dis. 2019, 80, S77–S84.

28. Wang, Y.; Holleufer, A.; Gad, H.H.; Hartmann, R. Length dependent activation of OAS proteins by dsRNA. Cytokine 2020, 126.

29. Donovan, J.; Dufner, M.; Korennykh, A. Structural basis for cytosolic double-stranded RNA surveillance by human oligoadenylate synthetase 1. Proc. Natl. Acad. Sci. U. S. A. 2013, 110, 1652–1657.

30. Schwartz, S.L.; Park, E.N.; Vachon, V.K.; Danzy, S.; Lowen, A.C.; Conn, G.L. Human OAS1 activation is highly dependent on both RNA sequence and context of activating RNA motifs. Nucleic Acids Res. 2020, 48, 7520–7531.

31. Hartmann, R.; Justesen, J.; Sarkar, S.; Sen, G.; Yee, V. Crystal Structure of the 2′-Specific and Double-Stranded RNA-Activated Interferon-Induced Antiviral Protein 2′-5′-Oligoadenylate Synthetase. Scand. J. Immunol. 2004, 59, 617–617.

32. Bignon, E.; Miclot, T.; Terenzi, A.; Barone, G.; Monari, A. Structure of the 5′ untranslated region in SARS-CoV-2 genome and its specific recognition by innate immune system via the human oligoadenylate synthase 1. Chem. Commun. 2022, 58, 2176–2179.

33. Miao, Z.; Tidu, A.; Eriani, G.; Martin, F. Secondary structure of the SARS-CoV-2 5’-UTR. RNA Biol. 2021, 18, 447–456.

34. Cao, C.; Cai, Z.; Xiao, X.; Rao, J.; Chen, J.; Hu, N.; Yang, M.; Xing, X.; Wang, Y.; Li, M.; et al. The architecture of the SARS-CoV-2 RNA genome inside virion. Nat. Commun. 2021, 12, 3917.

35. Lan, T.C.T.; Allan, M.F.; Malsick, L.E.; Woo, J.Z.; Zhu, C.; Zhang, F.; Khandwala, S.; Nyeo, S.S.Y.; Sun, Y.; Guo, J.U.; et al. Secondary structural ensembles of the SARS-CoV-2 RNA genome in infected cells. Nat. Commun. 2022, 13, 1128.

36. Tipo, J.; Gottipati, K.; Choi, K.H. High-resolution RNA tertiary structures in Zika virus stem-loop A for the development of inhibitory small molecules. Rna 2024, 30, 609–623.

37. Sharma, S.; Varani, G. NMR structure of Dengue West Nile viruses stem-loop B: A key cis-acting element for flavivirus replication. Biochem. Biophys. Res. Commun. 2020, 531, 522–527.

38. Fernández-Sanlés, A.; Ríos-Marco, P.; Romero-López, C.; Berzal-Herranz, A. Functional information stored in the conserved structural RNA domains of flavivirus genomes. Front. Microbiol. 2017, 8.

39. Bignon, E.; Marazzi, M.; Miclot, T.; Barone, G.; Monari, A. Specific Recognition of the 5′-Untranslated Region of West Nile Virus Genome by Human Innate Immune System. Viruses 2022, 14.

40. Waterhouse, A.; Bertoni, M.; Bienert, S.; Studer, G.; Tauriello, G.; Gumienny, R.; Heer, F.T.; De Beer, T.A.P.; Rempfer, C.; Bordoli, L.; et al. SWISS-MODEL: Homology modelling of protein structures and complexes. Nucleic Acids Res. 2018, 46, 296–303.

41. Case, D.A.; Betz, R.M.; Cerutti, D.S.; T.E. Cheatham, I.; Darden, T.A.; Duke, R.E.; Giese, T.J.; Gohlke, H.; Goetz, A.W.; Homeyer, N.; et al. No Title. AMBER 2016, Univ. California, San Fr.

42. Tian, C.; Kasavajhala, K.; Belfon, K.A.A.; Raguette, L.; Huang, H.; Migues, A.N.; Bickel, J.; Wang, Y.; Pincay, J.; Wu, Q.; et al. Ff19SB: Amino-Acid-Specific Protein Backbone Parameters Trained against Quantum Mechanics Energy Surfaces in Solution. J. Chem. Theory Comput. 2020, 16, 528–552.

43. Aytenfisu, A.H.; Spasic, A.; Grossfield, A.; Stern, H.A.; Mathews, D.H. Revised RNA Dihedral Parameters for the Amber Force Field Improve RNA Molecular Dynamics. J. Chem. Theory Comput. 2017, 13, 900–915.

44. Jorgensen, W.L.; Chandrasekhar, J.; Madura, J.D.; Impey, R.W.; Klein, M.L. Comparison of simple potential functions for simulating liquid water. J. Chem. Phys. 1983, 79, 926–935.

45. Mark, P.; Nilsson, L. Structure and dynamics of the TIP3P, SPC, and SPC/E water models at 298 K. J. Phys. Chem. A 2001, 105, 9954–9960.

46. Davidchack, R.L.; Handel, R.; Tretyakov, M. V. Langevin thermostat for rigid body dynamics. J. Chem. Phys. 2009, 130, 234101.

47. Feller, S.E.; Zhang, Y.; Pastor, R.W.; Brooks, B.R. Constant pressure molecular dynamics simulation: The Langevin piston method. J. Chem. Phys. 1995, 103, 4613–4621.

48. Hopkins, C.W.; Le Grand, S.; Walker, R.C.; Roitberg, A.E. Long-time-step molecular dynamics through hydrogen mass repartitioning. J. Chem. Theory Comput. 2015, 11, 1864–1874.

49. Miyamoto, S.; Kollman, P.A. Settle: An analytical version of the SHAKE and RATTLE algorithm for rigid water models. J. Comput. Chem. 1992, 13, 952–962.

50. Phillips, J.C.; Braun, R.; Wang, W.; Gumbart, J.; Tajkhorshid, E.; Villa, E.; Chipot, C.; Skeel, R.D.; Kalé, L.; Schulten, K. Scalable molecular dynamics with NAMD. J. Comput. Chem. 2005, 26, 1781–1802.

51. Phillips, J.C.; Hardy, D.J.; Maia, J.D.C.; Stone, J.E.; Ribeiro, J. V.; Bernardi, R.C.; Buch, R.; Fiorin, G.; Hénin, J.; Jiang, W.; et al. Scalable molecular dynamics on CPU and GPU architectures with NAMD. J. Chem. Phys. 2020, 153, 044130.

52. Humphrey, W.; Dalke, A.; Schulten, K. VMD: visual molecular dynamics. J. Mol. Graph. 1996, 14, 33–38.

53. Shitov, V. V.; Semenov, N.A.; Gozman, N.Y. PTRAJ and CPPTRAJ: Software for Processing and Analysis of Molecular Dynamics Trajectory Data. Telecommun. Radio Eng. (English Transl. Elektrosvyaz Radiotekhnika) 1984, 38–39, 14–16.

54. Fiorin, G.; Klein, M.L.; Hénin, J. Using collective variables to drive molecular dynamics simulations. Mol. Phys. 2013, 111, 3345–3362.

55. Ye, B.; Tian, W.; Wang, B.; Liang, J. CASTpFold: Computed Atlas of Surface Topography of the universe of protein Folds. Nucleic Acids Res. 2024, doi 10.1093/nar/gkae415.

