## Supplementary Information for "Molecular bases and specificity behind the activation of the immune system OAS/RNAse L pathway by viral RNA"

**Collective Variable Definition.**

To assess the global conformational changes related to the transition between the active and inactive form, we have used a collective variable based on the RMSD difference.
To this aim we have defined two reference structures composed of the OAS protein extracted from the HOLO (Active form) and the APO2 (Inactive form), respectively. At each step of the enhanced sampling simulation the RMSD of the OAS1 backbone has been calculated with respect to the Active ($RMSD_A$) and Inactive ($RMSD_I$) reference, respectively. The collective variable is calculated as:

(1) $\Delta RMSD = RMSD_I - RMSD_A$

As can be inferred from equation (1) when the system will be close to the Active reference the $\Delta$RMSD will be dominated by the $RMSD_I$ contribution, and hence will be positive. On the contrary conformations close to the inactive conformation will present negative values.

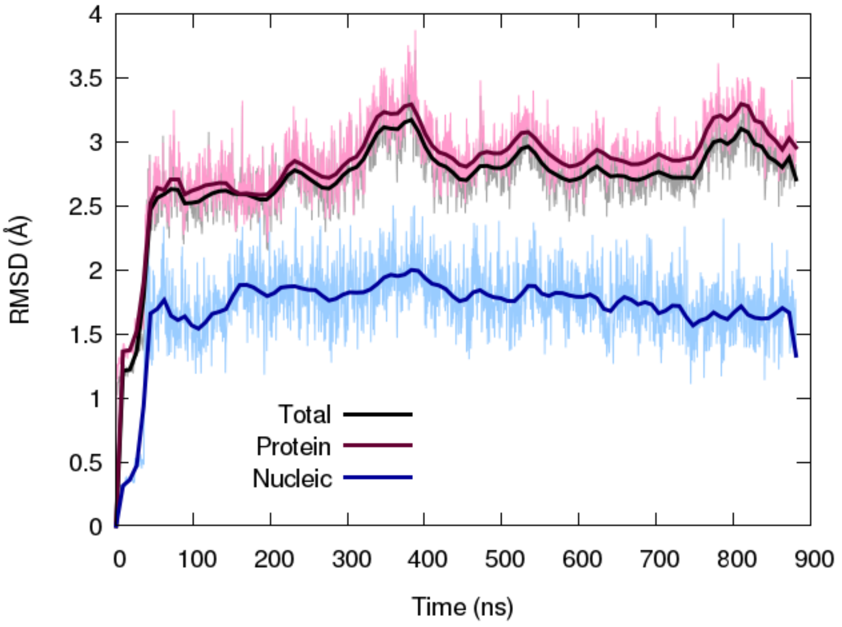

Figure S1. Time evolution of the RMSD for the HOLO system for the protein and RNA (Total, black), the protein (red), and the RNA (blue).

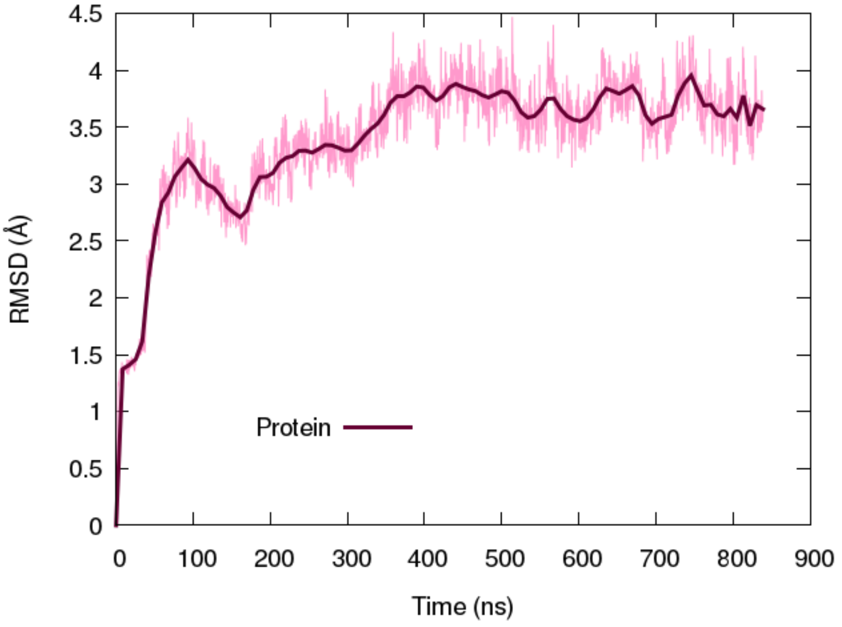

Figure S2. Time evolution of the RMSD for the APO1 system.

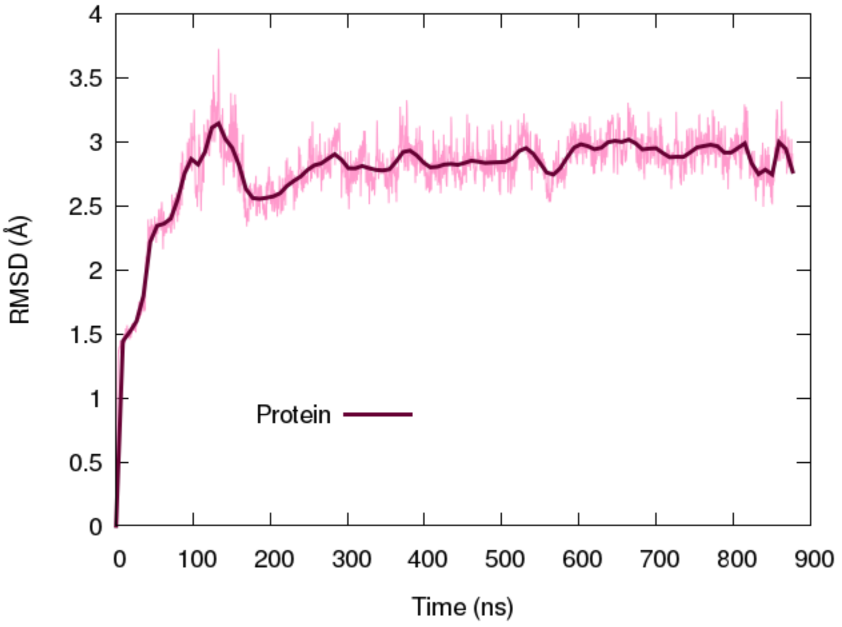


Figure S3. Time evolution of the RMSD for the APO2 system.

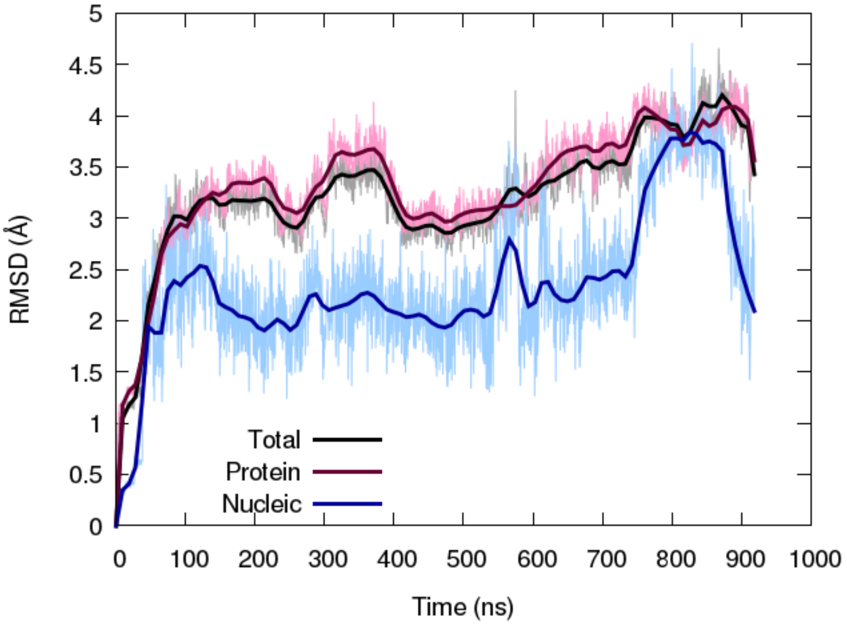


Figure S4. Time evolution of the RMSD for the APO2 system including RNA for the protein and RNA (Total, black), the protein (red), and the RNA (blue).

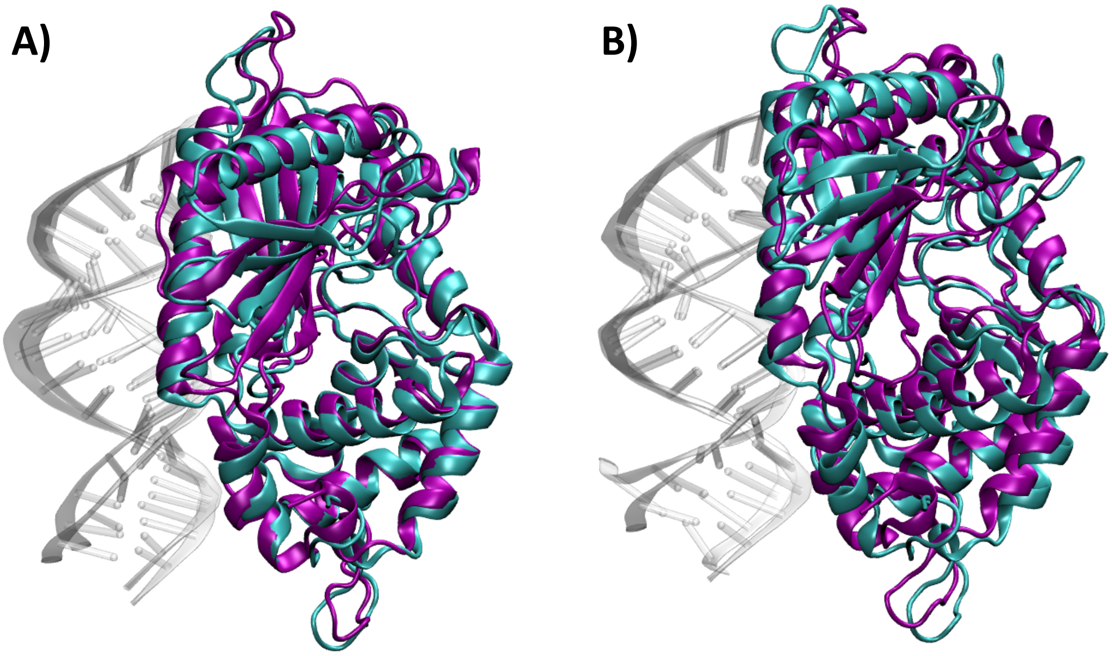


Figure S5. Superposition of the HOLO system (cyan) and the APO2+RNA system (purple) considering the most populated cluster (panel A) and the last frame of the dynamics (panel B).

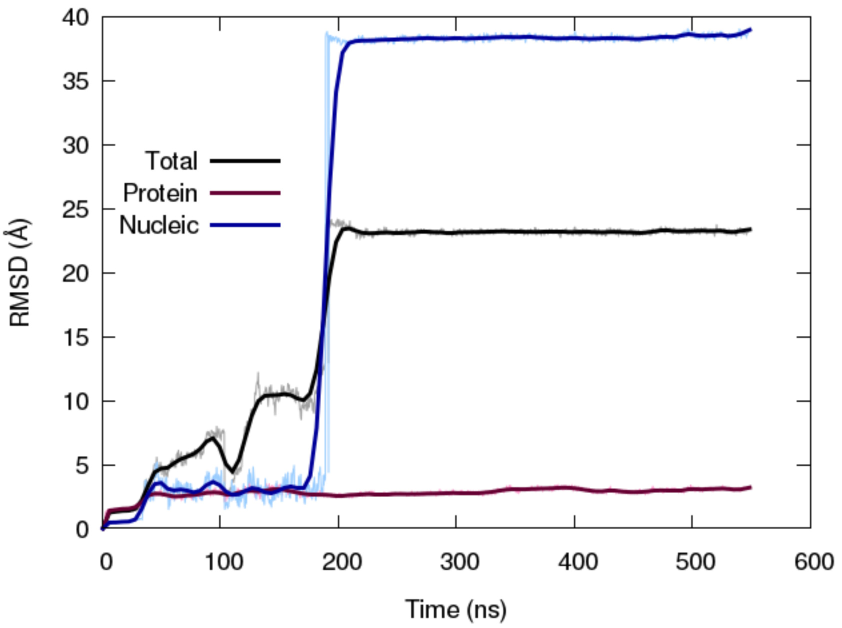


Figure S6. Time evolution of the RMSD for the HOLO17 system including RNA for the protein and RNA (Total, black), the protein (red), and the RNA (blue).

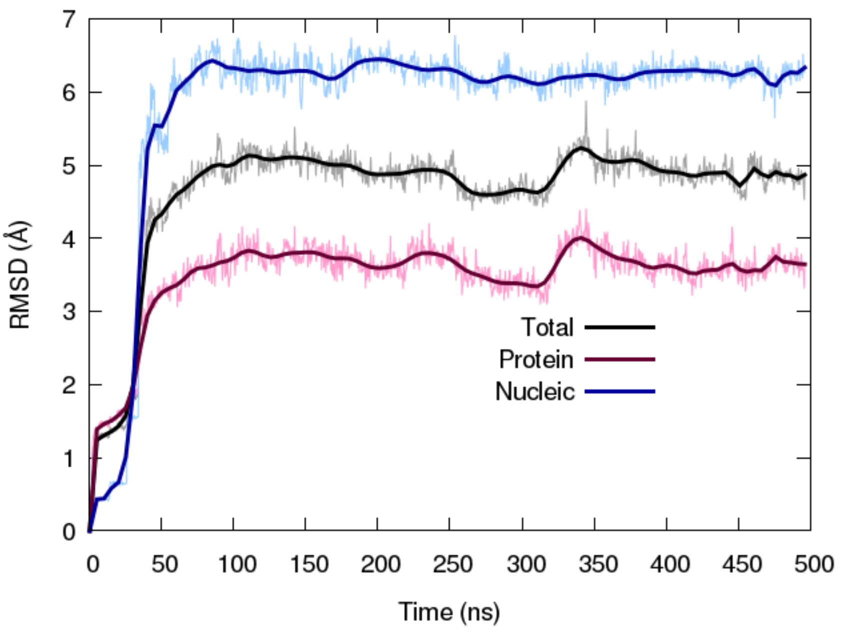


Figure S7. Time evolution of the RMSD for the HOLO18 system including RNA for the protein and RNA (Total, black), the protein (red), and the RNA (blue).

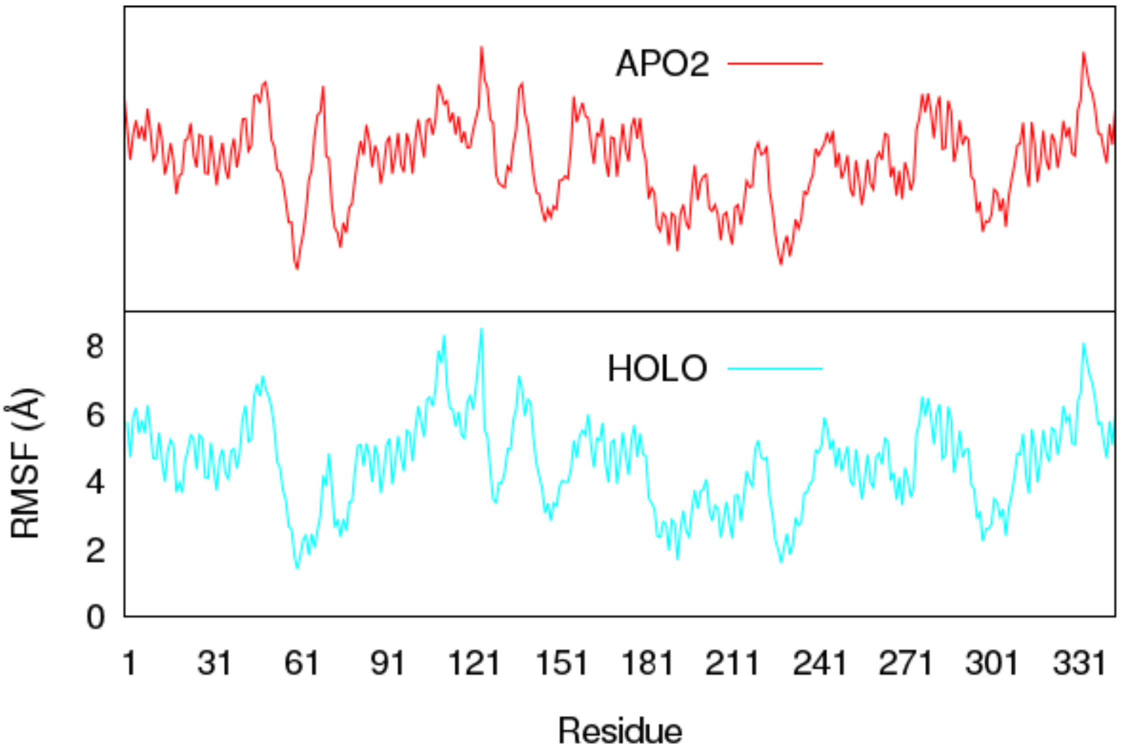


Figure S8. Per residue RMSF for the HOLO (bottom panel, cyan) and APO2 (top panel, red) systems.

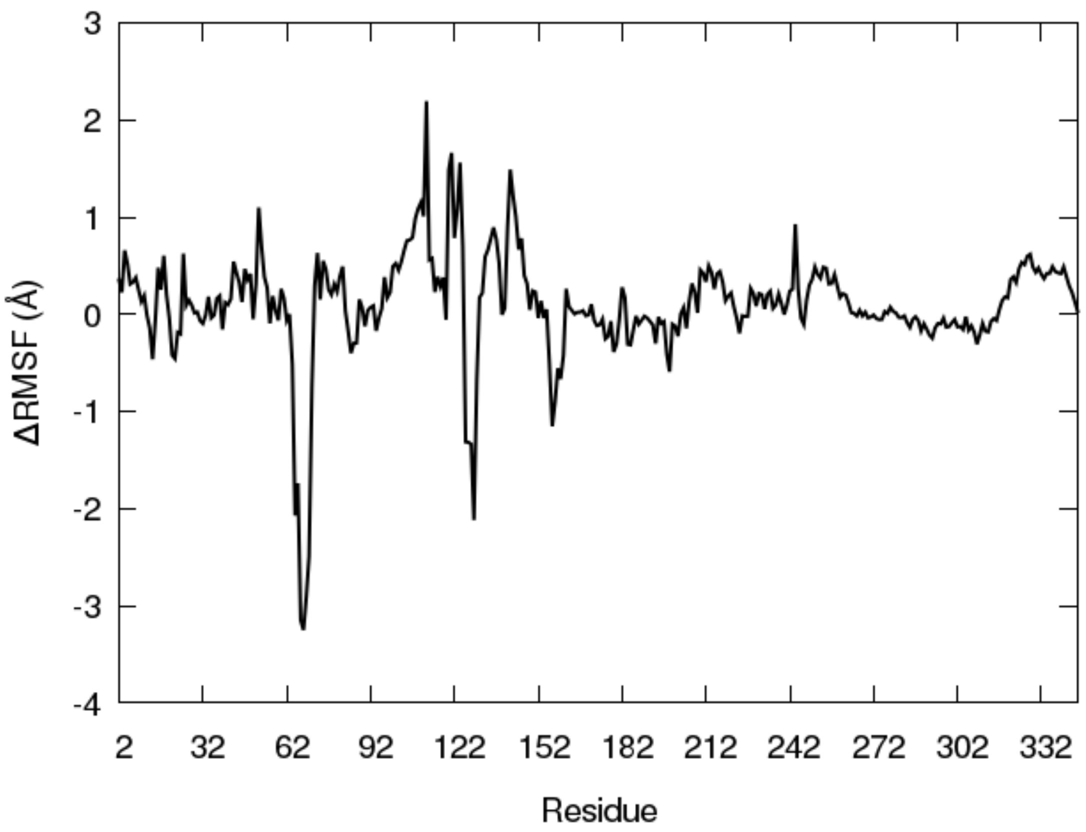


Figure S9. Difference of the RMSF (ΔRMSF) between the HOLO and APO2 systems.

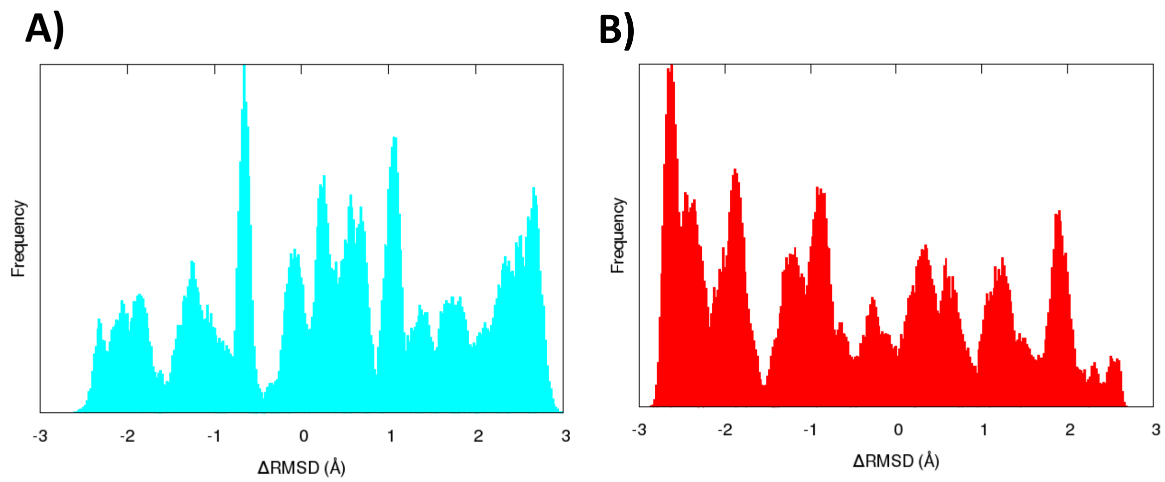


Figure S8. Distribution of the collective variable over the umbrella sampling for the HOLO (A, cyan) and APO (B, red) systems, showing the overlap of the different windows.
